# RAFTS: A graphical tool to guide Flux Simulator transcriptome simulation for method development in *de novo* transcriptome assembly from short reads

**DOI:** 10.1101/2022.07.13.499740

**Authors:** Matthew Doering, Jake M. Stout

**Affiliations:** Department of Biological Sciences, University of Manitoba, Winnipeg, Manitoba, Canada

## Abstract

Due to complexity of RNA transcripts expressed in any given cell or tissue, the assembly of *de novo* transcriptomes still represents a computational challenge when compared to genome assemblies. A number of modern transcriptome assembly algorithms have been developed to meet this challenge, and each of them have their own strengths and weaknesses dependent on the transcript abundance and complexity of the biological sample that is sequenced. As such, we are seeking to develop a transcriptome assembly pipeline in which multiple transcriptomes are generated, merged, and then redundancies are filtered out to produce a final transcriptome that should contain full length sequences of all transcripts. However, it is almost impossible to evaluate the efficacies of such novel assembly pipelines using short read sequencing data derived from biological samples due to not knowing *a priori* the transcript abundance and complexity. Thus, to test our pipelines we developed RAFTS. This tool is used to generate simulated short read sequencing datasets using annotated genomic data from model species.

## Introduction

The assembly of *de novo* transcriptomes, the puzzling together of short read RNA-Seq data to attempt to reconstruct the RNA present in a tissue or sample without the aid of a reference genome sequence, is a task with many approaches. A plethora of transcriptome assemblers are available (eg: Oases [1], TransABySS [2], IDBA-Tran [3], Trinity [4], STAble [5]), and decisions regarding the assembler algorithm, the impact of *k*-mer selection, and the use of read normalization can lead to different assemblies based on the same reads. Including methods that evaluate the quality of the assembly [6–8], or remove chimeras [9] and redundancy [10, 11] adds extra variables that further complicates the process. Recent recommendations for generating robust transcriptome assemblies are to use a variety of assemblers with different operating parameters, and then to evaluate the generated assemblies to select the most preferable assembly, or a set of assemblies that are subsequently merged [12]. It is currently unknown what combination of tools would yield the ‘best’ assembly using this approach, and there is thus a need to test a wide array of assemblers and post-assembly evaluation methods.

*In silico* RNA-Seq experiments can contribute to this by allowing for the generation of short reads with known genomic locations. This allows for *de novo* assembled contigs to be compared to be compared to the genomic sequences from which the reads were generated. For such *in silico* experiments there are various approaches that incorporate the simulation of library preparation methods to various extents, but the universal first step to generating simulated reads is to first simulate gene expression. There are various models for gene expression and transcript accumulation based on RNA-Seq data, the negative binomial distribution, gamma distributions, and other distributions. While these may be appropriate for particular types of transcriptomes and analyses, Flux Simulator is more broadly applicable to a wide variety of organisms, tissues, and transcriptomes because it uses a modified Zif’s law for simulation. Zipfian sets can arise when the underlying data may come from a variety of distributions.

The default parameters in Flux Simulator, k=-0.6, x0=9,500, x1=90,250,000, with 5×10^6^ transcripts accumulated, were set by comparison to mouse and human gene expression data. This also results in a fairly consistent number of genes being expressed in the simulation, which often needs to be varied based on the number of genes in the organism’s genome or number of genes either expected or desired to be expressed in the simulation.

To address this scenario and allow for a more complete and visual understanding of Flux Simulator’s parameter space for both gene expression and transcript accumulation, a total of 2,032,688 transcriptomes have been simulated across six species. The simulations show the consistency that Flux Simulator has across various genomes and replicated simulations from the human genome, visually summarized in the RAFTS html graphs, display the effect of parameter selection on the characteristics of the resulting transcriptome. Possible use cases are discussed.

Simulated short reads, *in silico* RNA-Seq experiments, in which the genome co-ordinates of each read are known, provide optimal reference assemblies that are useful for testing new assemblers [13, 14], comparing assembly methods [12, 15], developing of methods for read-alignment [16, 17], and the training of computational models [18]. While numerous read simulation software options are available for simulate sort-read genome sequences [19], fewer options are available for simulating RNA-Seq reads [20] (Table 1). Some genome-based read simulators may also be appropriate for RNA-Seq simulations if the input can be a set of fasta sequences, which allows for the use of transcripts for RNA-Seq simulation as in the case of dwgsim. However, RNA-Seq read simulators generally allow for alternative splicing and other types of RNA editing as well as finer control over simulating experimental methods themselves.

Of the RNA-Seq read simulators, Flux Simulator [21], rlsim [22], and Polyester incorporate experimental method simulations. Only Flux Simulator, the first algorithm for realistic RNA-Seq read simulation to have been published, can simulate all the experimental method, including sequencing library preparation steps. It allows for simulated control over expression levels, transcription start site variability, polyadenylation, fragmentation, reverse transcription, size filtering of fragments, amplification including settings for GC amplification bias, and sequencing with error models.

Like Flux Simulator, rlsim introduces experimental biases into the sequencing results. It has parameters for polyadenylation, fragmentation and hexamer priming, PCR amplification, and size selection, specifically focusing on the PCR and size selection steps with tools to evaluate GC-dependent amplification efficiency and insert size distribution. Polyester, which provides fragmentation and sequencing parameter sets, GC bias models, and sequencing error models, specifically addresses replication and differential expression to allow for fine scale control over read counts from each transcript for each simulated sample library [23].

RSEM (RNA-Seq by Expectation Maximization) [24], SimSeq [25], and RNA-Seq Simulator (popmodels.cancercontrol.cancer.gov/gsr/packages/rna-seq-simulator) learn experimental parameters from an experimentally acquired RNA-Seq dataset, and then applies that to produce simulated reads. SimSeq and RNA-Seq Simulator only extract levels of gene expression from the input dataset. RSEM additionally estimates fragment length distributions and error models. RNA-Seq Simulator uses gene expression levels to simulate datasets with differential expression or overdispersion characteristics.

TuxSim [26], like Polyester, is designed for replication with built in methods for the perturbation of gene expression in set ways for controlled up or down regulation of some or all isoforms of a gene. RNASeqReadSimulator (alumni.cs.ucr.edu/~liw/rnaseqreadsimulator.html), another unpublished simulation option, allows for control over error rates and positional biases. BEERS (Benchmarker for Evaluating the Effectiveness of RNA-Seq Software) [27] was created to evaluate the alignment of reads to a reference genome, not specifically for *de novo* methods, however it allows for control over the source of polymorphisms in reads, including partial retention of introns and possible insertions or deletions [28].

In addition to simulating experimental methods, sequencing errors and models for gene expression and transcript accumulation (i.e. read counts per transcript) underpin each simulation method. The resulting simulated reads depend to a great extent on the assumptions underlying the simulated gene expression. This is analogous to RNA-Seq reads being impacted by the underlying transcript accumulation levels; more frequent reads and *k*-mers can be indicative of higher transcript accumulation. Both Polyester and TuxSim use the negative binomial distribution, though Polyester uses the distribution to simulate read counts whereas TuxSim uses it to simulate the number of fragments accumulated from each gene. The negative binomial model, which is also known as the gamma-Poisson model [29, 30] and used by Polyester and TuxSim, has been shown to be appropriate for modelling transcripts accumulating from expressed genes [31–33]. The negative binomial model has two parameters, and approaches the Poisson distribution, with a single parameter λ, as the mean, μ, and variance, σ^2^, converge (Figure 1). The negative binomial model is a robust alternative to the Poisson for modelling count data with events being positively correlated [29, 34].

**Figure 1.**
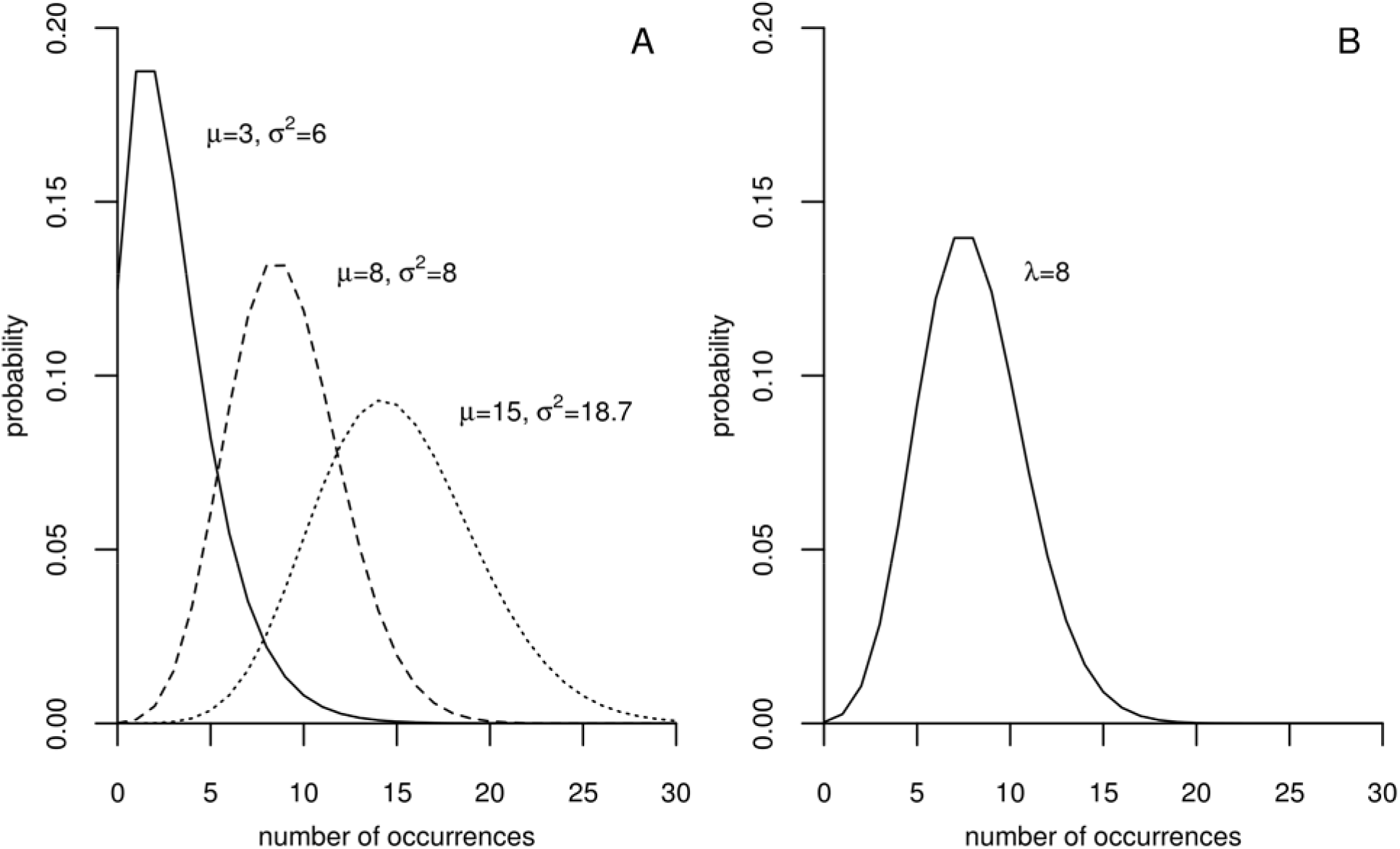
Density plots of three negative binomial distributions (A) and a Poisson distribution (B).

Rlsim, on the other hand, models expression levels for each gene based on mixed gamma distributions. The gamma distribution models wait times till the *k*^th^ Poisson event. The gamma distribution itself, though perhaps useful in some particular scenarios such as chromatin remodelling [35, 36] or when the assumptions or necessary simplicity are warranted [37], has been shown to be unlikely to be generally indicative of biological gene expression data [37, 38]. Rather than modelling mRNA accumulation, gamma models may be more appropriate for modelling levels of some cellular proteins, especially protein species with fewer than 10 molecules per cell [39], with the two gamma parameters, *a* and *b*, corresponding to transcription rate and the duration of stochastic translation bursts [40], respectively.

Accounting for repressor activity in transcription results in log-normal gene expression [40], which is the model used by RNASeqReadSimulator to model numbers of transcripts accumulating from expressed genes. In comparison to a normal distribution, log-normal modelling of gene expression has been shown to be appropriate for some genes [41]. The implication of log-normal transcription, as discussed by Bengtsson et al. [41] is that gene expression measured for a collection of cells is not reducible to a single-cell scale. There is some indication, however, that not all genes follow the same expression profile with many different models being appropriate for different subsets of expressed genes [42]. And, in cancer cells, changes in the distribution of gene expression can be as informative as fold changes in gene expression [42]. Therefore, the aim of simulating a biologically realistic dataset, without relying on previously published RNA-Seq reads, becomes a complex task.

Flux simulator [43] takes a different approach by modelling transcript accumulation after Zipf’s law [44]. This allows for the accumulation of more rare transcripts than other distributions, including negative binomial distribution. Griebel et al. developed the Flux simulator based on the observation that most expressed sequence tags are rare and few are very abundant [45]. The greater contribution at the time was the level of control over the simulated experimental protocol to more realistically generate simulated reads accounting for experimental biases. With the emergence of additional RNA-Seq read simulators, a greater review of Zipf’s law and its applicability to transcriptome simulation is useful.

Zipf’s frequency distribution arose in a linguistics context, and was later generalized in the same context by Mandelbrot [46]. This distribution shows an inverse relationship between frequency and rank that is linear on a log-log scale,

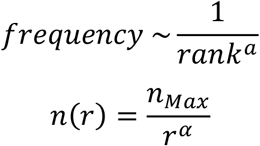

In linguistics *α* generally equals 1, and it is this form that is typically referred to as Zipf’s law. In the second form, *n* is the number of occurrences, *r* is rank, and *n*_*Max*_ is the maximum number of occurrences in the set. The result of this is that there are few observations (i.e. ranked items) with high frequency and many items with low frequency.

Zipf’s law, also known as a Pareto distribution, is one example of a power law. Power laws and Zipfian distributions have been shown to occur throughout human language and society as well through the natural world. In addition to occurring in many human languages [47], Zipf’s law has been observed in computer software and programming languages [48, 49]; in music [50], including in aspects such as melodic intervals, loudness, and pitch [51]; an in such cognitive processes as memory and perception [52]. The societal prevalence of power laws is apparent through the number of deaths during wars [53], city sizes [54], and national GDPs [55]. In the natural world, phenomena as disparate as the energy levels of earthquakes [56], asteroid sizes [57], and the size of solar flares [58] all follow power laws.

Given this range of areas in which power laws can be found, the question as to why this this is so comes to the fore. Numerous explanations, some of which are briefly described below, have been proposed for various scenarios in the century since Zipf first described his power law for word frequencies.

### Combination of distributions

In a linguistics context, Zipfian word frequencies from random typing have been interpreted as an exponential function of words with length *y* with the total possible number of words having length *y* also being exponentially distributed [59, 60]. Another example of this combination of distributions has been proposed to explain gene family and genera sizes which may be seen to rely on exponential growth and random decay functions [61]. Reed and Hughes [61] also demonstrated, using a Poisson growth function with decay expressed by the Galton-Watson probability of name extinction [62], that the Zipfian nature of surname distributions can also arise through a combination of distributions.

### Yule process

Yule [63] described a process to explain the power law distribution of the number of species within taxonomic groups. According to this model, assume a set of categories with new categories being added occasionally. Between the addition of categories, individuals are added to various categories with a probability proportional to the size of each category, i.e. larger categories are more likely to gain additional members and gain those members at faster rates than smaller categories. An alternative explanation of it is that if the appearance of categories during a time period and the addition of individuals are both described by Poisson or negative binomial distributions then this leads to the membership sizes of the categories having a power law distribution [64, 65]. Because Yule did not account for extinction of species, this process does not realistically represent species distributions as intended. However, it has gained acceptance in describing, for example, paper citations [66] and web page links [67].

### Criticality and self-organized criticality

Given long time or spatial scales governing some systems, power laws may arise at the critical points of phase transitions at some points along those scales [68, 69]. In this case the appearance of power laws is a unique characteristic of very particular sets of conditions that may sometimes exist.

Seeking to explain the apparent noise in power laws arising in temporal scenarios, Bak et al. [70] proposed that some systems, in the self-organized criticality (SOC) model, may converge on and fluctuate around a critical point at which a power law exists, with power laws not describing the data at parameter values far away from that critical point. With systems, as diverse as the flow of the Nile River and the luminosity of stars, fluctuating around such “minimally stable” states, power laws can appear to be prevalent across many different systems.

### Highly optimized tolerance

The highly optimized tolerance (HOT) model [71] describes how power law distributions arise from scenarios that are optimized to balance trade offs between yield, resource cost, and risk tolerance. These tend to be highly efficient systems that are able to withstand the uncertainty of chance and rare events [72] and power law distributions tend to quickly emerge with even a small amount of organization applied. HOT, in contrast to criticality models which position systems at boundaries of order and chaos, use structure and redundancy to perform reliably in uncertain environments [71].

### Coherent noise

First proposed to model avalanches and species extinction [73], the coherent noise model assumes larger scale, global, external “noise” forces driving organization toward power law distributions. This is different than the SOC models in that this noise is coming from variables not accounted for by SOC model parameters [74]. Additional models leading to Zipfian and power law distributions include fragmentation processes [75], random multiplicative processes [76], inverses of quantities [77], random walks [78], and random extremal processes [79].

A Zipfian set is a whole, resulting from neither a subsetting nor a merging of sets [55]. It cannot be an assemblage so as to maintain the ordering of items because the order of items will change when sets are combined. It has been speculated, though not demonstrated, that deviation from Zipf’s law can show a breakdown in structure, or that multiple processes at work.

In the particular case of gene expression, a strictly Zipfian model can begin to break down with infrequently expressed genes and lowly accumulated transcripts departing from the expected frequencies. For this reason an exponential decay factor is incorporated into Flux Simulator [21]. Such a model has been termed a “weak” inverse power law [80].

Zipfian distributions have been shown to describe transcript accumulation in human, rat, mouse, yeast, nematode tissues, as well as human caner cell lines, *Arabidopsis*, and *Pinus* [45, 81, 82]. To explain the presence of this distribution, thermodynamic and evolutionary models have been proposed. The evolutionary model assumes that DNA mutations in cis-regulatory elements lead to random variation in gene expression levels and that gene duplication leads to gene number increasing over generations [81]. The thermodynamic model is based on the assumption of the existence of a critical state defined by a maximal cell size with the chemical similarity to “parent” cells maximized following cell division [45].

The usefulness of using a Zipfian distribution for the basis of transcriptome simulation may be its approximation of underlying behaviour rather than understanding it as an emergent, universal property. The rationales for Zipfian and power law distribution use may be as diverse as the fields of study in which they are used. In the case of simulating transcriptomes, using a Zipfian distribution can be seen as a useful model to capture a great deal of biological complexity without the necessity of directly modelling that complexity.

This frequency distribution is appropriate for transcriptome assembly applications where more than half of the total mRNA is accumulated from the expression of less than 100 genes, and the expression of thousands of genes make up the remainder of the mRNA pool [83]. This model has been shown to be appropriate to a variety of eukaryote transcriptomes [43, 45, 81, 82]. In diverse contexts where the observed frequencies span many orders of magnitude, as in copies of mRNA in transcriptomes, distributions in line with Zipf’s law become prevalent [84]. Aitchison et al. [84] showed that a broad, Zipfian distribution can be assembled, among other ways, from a set of narrow, non-Zipfian distributions. As discussed above, a variety of distributions, each applying to a set of genes and expression levels, taken together is necessary to understand the expression profile of a transcriptome. Therefore Zipf’s law can provide a transcriptome simulation method to capture the biological complexity of the various gene expression distributions present in a transcriptome without the need to account for each distribution individually.

Flux simulator allows control over expression levels with a set of parameters and produces reads genome-annotated transcripts. It additionally offers control over all simulated steps of expression, library preparation, sequencing. Introduces experimental biases into the simulated reads. Although Flux Simulator doesn’t explicitly allow for control over the number of genes expressed, the number of RNA molecules accumulated is a user-controlled input parameter. Flux Simulator allows for control over the exponent *k* of Zipf’s law, in the range of 0 > *k* > -1, and also provides an additional exponential decay component to reduce transcript accumulation levels with large numbers of genes.

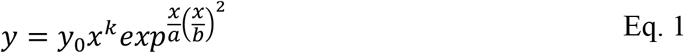

Equation 1 is given by Griebel et al., though, the result of an apparent typo, lacks the required negative sign in the exponent of the exponential decay component. Here *a* is *x*_*0*_ and *b*^*2*^ is *x*_*1*_. This can be rewritten to more clearly match the parameters provided to Flux Simulator as equation 2. The value of *y*_*0*_ is the maximum number of transcripts accumulated from the most highly expressed gene (10^4^ for the mammalian datasets investigated by Griebel et al.)

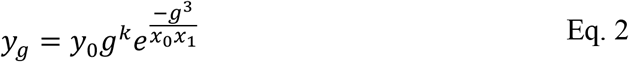

Here *y* is the number of transcripts accumulated from gene *g* and *k, x*_*0*_, and *x*_*1*_ are the values provided for those variables.

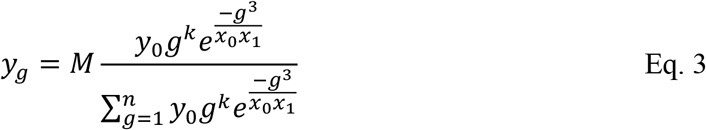

To more completely express the Flux Simulator method, equation 3 normalizes number of transcripts accumulating from gene *g* by dividing by the sum of all transcripts accumulated. The normalized number of transcripts sums to 1 across all expressed genes. Multiplying by *M*, the number of molecules desired to be simulated and given by NBMOLECULES in Flux Simulator, results in the scaled number of transcripts accumulated from gene *g*.

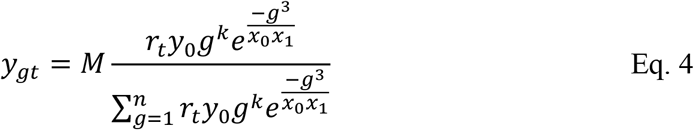

Given that there is variation across Flux Simulator transcriptome simulations with constant values of *k, x*_*0*_, and *x*_*1*_, there is an element of randomization incorporated into the simulator. In equation 4 it is expressed as *r* for the *t*^*th*^ transcriptome simulated. When *k*=0, the simulator expresses all annotated genes with a mean expression level of 15 and a standard deviation of 5 [43].

The default values used by Flux Simulator are *k* = -0.6, 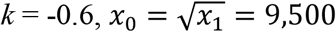, NBMOLECULES=5×10^6^. While these are appropriate for some instances, they result in the simulated expression of approximately 19,000 genes. Depending on the organism and tissue for which a simulated transcriptome is sought, this may not lead to an accurately modeled dataset due to an unrealistically high or low number of genes represented.

By using real datasets, methods like RSEM, SimSeq, and RNA-Seq Simulator may be seen to maintain the underlying complexity of RNA-Seq data by making no assumptions of the underlying levels of transcript abundance, as opposed to the assignment of levels of gene expression or read counts from parametric distributions. By using Zipf’s law with a degree of randomization, Flux Simulator’s method is something of a compromise between maintaining the complexity of experimentally-acquired RNA-Seq data and the assignment of gene expression probabilities by parametric methods. Thus, Flux Simulator appears to be the best option to model a transcriptome without the introducing errors resulting from a lack of familiarity with a particular dataset. The use of multiple simulators has been employed to control for simulation biases arising from a single simulator [85].

While possible to use the Flux Simulator equation to select values for NBMOLECULES, k, x_0_, and x_1_ to give expression profiles with desired characteristics for a targeted number of genes to be expressed, it is not a task readily accomplished. To meet that need, we present a graphical tool to select parameters to control gene expression simulations (both numbers of genes expressed and of transcripts accumulated in broad categories of expression). This tool allows Flux Simulator to be used across organisms to simulate the expression of varying numbers of genes and levels of expression to tailor simulated RNA-Seq reads to particular experimental requirements. The resulting levels of transcript accumulation can be used to simulate RNA-Seq reads using Flux Simulator or can be used as input for a different RNA-Seq read simulator.

Online versions of the Flux Simulator parameter selection graphs are available at home.cc.umanitoba.ca/~umdoeri0/flux-simulator.

## Methods

Zipfian distributions may arise through several means and have been shown to model the complexity of mRNA accumulation across many organisms. Because of this Flux Simulator, which is based on a Zipfian model, is an appropriate tool to simulate gene expression, the first step in simulating short reads necessary for bioinformatic method development and tool evaluation. Flux Simulator relies on several parameters to control the number of genes expressed and the levels of transcript accumulation, but lacks a user-friendly method to select the parameters necessary to tailor the simulated transcriptome to a particular organism or tissue. To fill this need, we have investigated the Flux Simulator parameter space and developed the RAFTS tool to allow for a visual understanding of Flux Simulator’s transcriptome simulation parameters.

A first set of simulations was created to span a wide range of parameter values in order to begin investigating the numbers of genes expressed by Flux Simulator, using the current *Brassica napus* genome assembly (Table 2). The parameters used spanned several orders of magnitude above and below default values for number of molecules, x_0_, and x_1_. For *k*, our selected values span the full range from 0 to -1, excluding the end values. (Supplementary Table 1). With a single replicate for each simulation, this resulted in 146,608 sets of parameters.

The resulting parameter files were parsed to count the number of genes expressed, the number of transcripts accumulated, and the numbers of genes expressing transcripts that accumulated to super-prevalent (SP), intermediate (I), and complex/rare (CR) levels. While these thresholds vary in the literature, the thresholds we report are consistent with those used by Flux Simulator with SP transcripts having > 500 copies and CR transcripts having fewer than 15 copies. Martin and Pardee [86] used these thresholds, saying that abundant transcripts had > 500 copies and middle abundant transcripts had 15-500 copies per cell. They added an additional category of rare transcripts being those with < 5 copies per cell, which represents the largest portion of transcripts with reports of up to 86% of all transcripts belonging to this category [87]. Mortazavi et al. [88] defined rare transcripts as having 10 or fewer copies per cell. Williams [89], in contrast, used thresholds of 14-500 for middle abundant transcripts. Less than 0.5% of genes expressed tend to accumulate transcripts to a super-prevalent level [87]. This represents fewer than 100 genes [87], and can be as few as 5-10 genes [90].

We used the SP, I, CR thresholds in two ways: raw and scaled. In the raw summaries, we used a straight count of the number of transcripts accumulated from each gene for categorization. With the scaled summaries, we divided the raw count by the total number of mRNA molecules simulated and multiplied by 100,000, using the same assumption of 100,000 transcripts per cell as the simulator itself.

These summaries were computed with an in-house bash script and the results written to a csv file. We then plotted the summarized data using the Data Driven Documents (D3) JavaScript library v. 4 [91]. We plotted, in separate graphs across number of molecules accumulated, numbers and percents of genes expressed in total and in each of the three expression categories, and numbers and percents of transcripts accumulated in total and in each of the three categories. The curveMonotoneX function was used to plot the lines. This ensures that the lines pass through each data point, and does not allow for local maxima or minima to exist between data points [92]. Drop down menus with the full range of parameter values simulated for k, x_0_, and x_1_ were implemented to dynamically update the number of genes or transcripts in each category. On hovering over the graphs and moving the cursor position, the co-ordinates (number of molecules in the simulation, number of genes/transcripts) are reported on the graph, interpolated according to the D3 graphing function used. As the cursor location is reported as a percentage of the line length, not all reference markers move similarly and so attention to x-values across the lines is necessary.

This preliminary, version 1, set of graphs was used to extract sets of parameters to simulate biologically realistic, in terms of numbers of genes and SP transcripts arising from the expression of fewer than 10 genes, transcriptomes in *Brassica napus, Arabidopsis thaliana, Mus musculus, Saccharomyces cerevisiae*, and *Homo sapiens*. Current genome assembly and annotation versions were used for each species (Table 2). Based on numbers of genes found to be expressed in various tissues in each species, we decided to select parameters for the simulation of approximately 34,000 genes in *B. napus*, approximately the number expressed in germinating roots [93], ∼10,000 genes in *A. thaliana*, similar to the number expressed in a germinating seed or in the hypocotyl or root [94], ∼8,000 in *M. musculus*, similar to the number expressed in brown fat, thymus, and brain [95], ∼25,000 in *S. salar*, between the numbers expressed in the liver and expressed in the ovary [96], ∼4,500 in *S. ceresvisiae* [97], and ∼15,500 in *H. sapiens*, similar to the number expressed in the testes [98] and different from the other expression scenarios already created. Default transcriptome simulation settings were also used and, based on the version 1 graphs, estimated to express 18,800 genes. All transcripts present in the genome annotations were used as input to Flux Simulator with two exceptions: only representative gene models were used in *A. thaliana* and only the genome features annotated as the best reference sequences were retained in the *M. musculus* annotation.

Using these 7 sets of parameters (Table 3), 10,000 replicates with each set for each species’ genome annotation were performed. The gene expression and transcript accumulation characteristics of the resulting transcriptome models were summarized (Table 4, 5).

Because there is variation in the simulation method used by Flux Simulator, replication is needed in the web-based graphical summary for there to be confidence in using it as a predictive tool for transcriptome simulation. We decided to use the *H. sapiens* genome for this version 2 of the summary graphs because it had the largest number of transcripts and splicing variants recognized by Flux Simulator of the analyzed species (112,981; Table 3). Based on those results, we calculated the margin of error *m* expected in the raw and scaled numbers of genes expressed and transcripts accumulated assuming 10 replicates, 95% confidence intervals, and the means and standard deviations calculated for each parameter set for *H. sapiens* (Table 6).

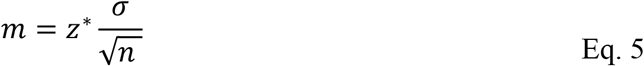

Given these assumptions, the number of genes expressed and the number of genes producing CR transcripts are accurate to within 2% of the mean, and the number of genes producing I transcripts is accurate to within 10% of the mean. The larger variations in genes producing SP transcripts is due to the much smaller number of genes expressed in this category. But where the margin of error appears inflated as a percent, it is a single digit number of SP genes. Ten replicates were done to balance the computational intensity (requiring approximately 70 days when run with 24 threads and 256 GB RAM) with reliable estimation of the variation within each combination of parameters evaluated.

Using the same parameter sets (Supplementary Table 1) as for the version 1 graphical summary, we did 10 replicate transcriptome simulations from the *H. sapiens* genome with Flux Simulator and used the averages to create version 2 graphs.

All simulations were run on a server with 48 processors and 512 GB RAM running Ubuntu 16.04.6 LTS. Each instance of Flux Simulator was run using 21 GB memory.

## Results

We investigated the parameter space of Flux Simulator’s transcriptome simulation method. In addition to developing the RAFT graphical tool, a more complete understanding of the underlying simulation method can be gained.

Because the different replicates produced slightly different gene expression profiles, there is a randomization component to the implementation of Zipf’s law used in Flux Simulator. If there were no element of randomization in the algorithm, then every replicate simulation with the same set of parameters would be exactly the same with the only variation coming from the random ordering of genes.

The other insight that can be gained into Flux Simulator transcriptome simulation is that not all gene regions annotated are recognized by the simulation engine. In the case of the *B. napus* genome more than half of the annotated gene model variants present were not included in any of the model transcriptomes. This is not due to splicing variants not being recognized by Flux Simulator, as 111 representative gene models were not recognized (Table 7). This draws attention to the importance of preliminary test simulations to ensure the usability of the annotations when the simulation experiment requires the availability of particular gene models for expression and subsequently sequencing.

The number of genes for which expression is simulated remains constant across simulations, provided there are sufficient gene models in the annotation (Table 4). As would be expected drawing from Zipf’s law, the first genes to not be expressed when too few models are available are those with CR expression profiles. This is evident in simulation set A with *S. cerevisiae* (Table 4, 5). In that situation, more genes with SP and I expression profiles are expressed in order to maintain the set number of transcripts requested in the simulation. Similarly, the genes with SP expression profiles are always expressed and are used to inflate the number of transcripts accumulated (Table 4, simulation set D).

Using the defaults parameters gives realistic transcript accumulation distributions, but not realistic numbers of genes expressed. In many cases expressing nearly 19,000 genes would represent an RNA extraction from multiple tissues. The desirability of this depends on the purpose of the transcriptome and read simulations. If the goal is a biologically realistic number of genes from a particular tissue is desired, then the mark would be missed. An additional concern would arise if the annotation used for simulation contained fewer than 19,000 gene models, in which case the expression profile would not be maintained and some or many expected rare transcripts would be absent from the simulated transcriptome and sequenced reads.

If the purpose of the simulation is to simulate expression of a known set of genes with an expression profile following Zipf’s law, then selecting parameters to simulate expression of that known number of genes with the desired expression profile is possible. We did this with the parameter selection in simulation set A with the *M. musculus* genome (Table 4, 5). The expectation was to simulate the expression of approximately the same number of genes as were in the genome annotation used and in fact all the genes in all the simulations were expressed. Both the numbers of raw and scaled genes expressed and transcripts accumulated were fairly consistent with the simulated expression profiles from genomes with more gene models.

Increasing the number of molecules accumulated in the transcriptome simulation is akin to the amount of tissue used for RNA extraction. More transcripts are recovered, and more that accumulate to very low levels. In terms of parameter selection, our experience in selecting the parameter sets used for each species mirrors that of Griebel et al. [43] with a suitable range for values of k being from -0.6 to -0.9 for biologically realistic expression profiles.

**Figure 1.**
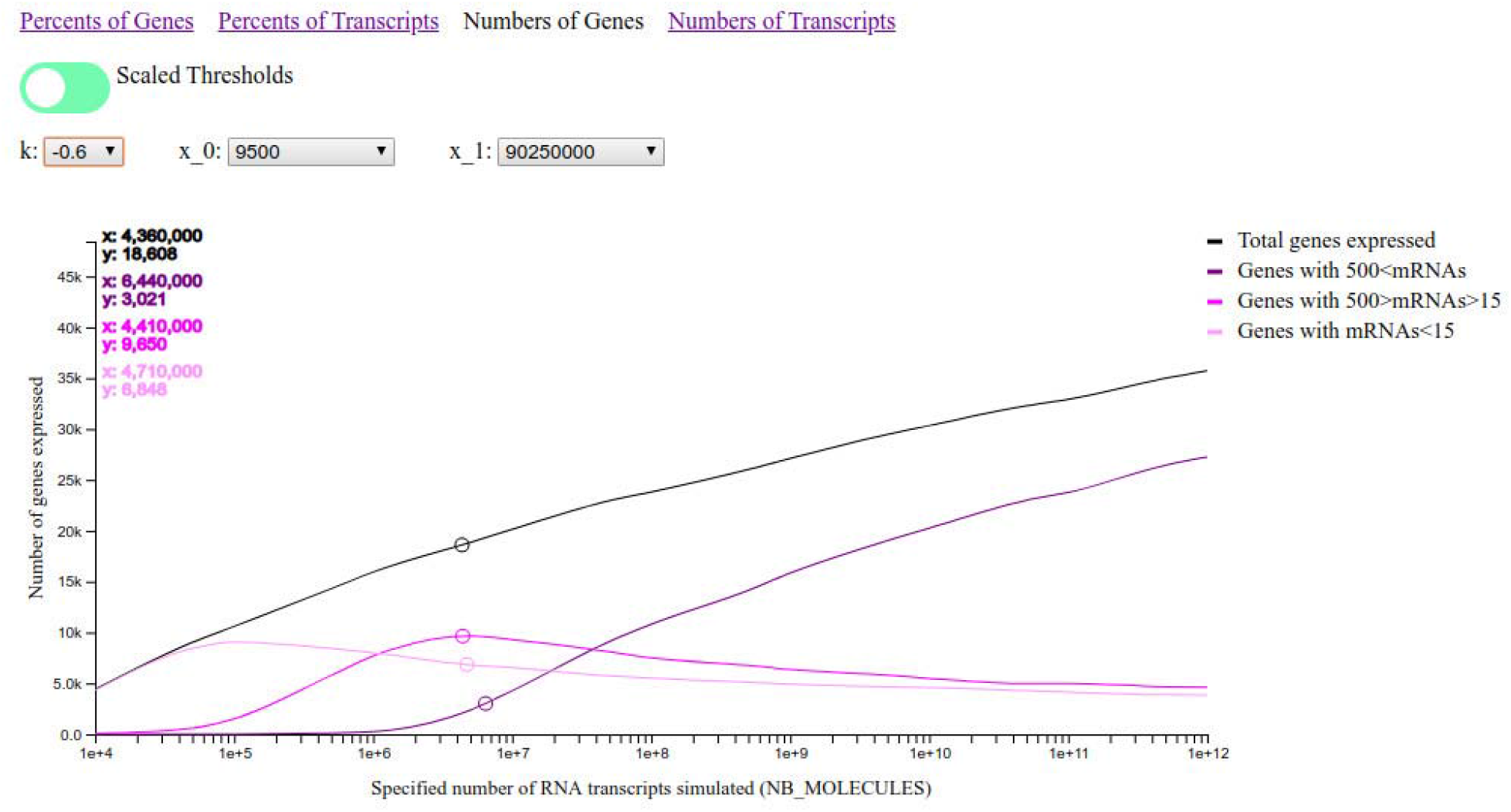
Screen capture of the RAFTS graphical tool to summarize the numbers of genes expressed and transcripts accumulated as Flux Simulator parameters vary. The open circles move with the cursor are showing approximate default values (5×10^6^) for the number of mRNA molecules accumulated.

## Discussion

We presented a graphical simulation model, RAFTS, to inform parameter selection for transcriptome simulation, the first step of short read simulation, with Flux Simulator. While RAFTS is not a novel read simulation method, it is an extension of the scope for which the common read simulator Flux Simulator can be used. Three use cases are presented below:

1. read simulation and how this RAFTS can prove useful in simulating reads from simulated transcriptomes with biologically-informed characteristics,
2. simulating reads from simulated transcriptomes with particular expression profiles, while maintaining Zipf’s law,
3. simulating transcriptomes, with calculated read counts for each gene, with expression profiles based on Zipf’s law, rather than parametric distributions, to use as input into other read simulators for particular replication or differential expression purposes.

### 1. Simulate biologically-meaningful transcriptomes to sequence

Depending upon the species, greater flexibility in defining the number of genes expressed, not only the total number of mRNA molecules accumulated, can be valuable. For instance, using the transcriptome simulator’s default settings simulates the expression of 18,952 genes, on average, but this is probably twice as many genes as would normally be expressed in any given *Arabidopsis* or mouse tissue and certainly far more than the number of genes even present in a yeast genome. Using the graphical tool we present here, parameters can be selected to allow for gene expression to follow Zipf’s law while expressing a realistic number of genes. Carrying such a simulated transcriptome through Flux Simulator’s library preparation and sequencing simulations will result in greater control over the biological relevance of the resulting reads.

Though other read simulation tools allow for experimentally sequenced RNA-Seq short read datasets to be used to inform the simulation parameters, the precise experimental details, protocols, and sources of error are not necessarily known when acquiring a publicly available dataset. Therefore, to avoid simulating an undesirable artefact in the dataset, the value of simulating gene expression and library preparation becomes more apparent.

### 2. Simulate transcriptomes with particular characteristics

While simulated reads resembling experimentally acquired data, with the benefit of possessing known genome co-ordinates, can be valuable, equally valuable can be controlling the simulation of gene expression more closely to allow for only the sequencing of a particular category of transcripts. For instance, parameters could be selected to allow for the expression of 10,000 genes all of which accumulate fewer than 15 transcripts. Such an approach, though not biologically accurate, would allow for *de novo* method development, including for *de novo* transcriptome assembly, to more precisely target and evaluate performance on solving particular problems and identifying unique methodological strengths and weaknesses. Expanding that concept, multiple simulated read datasets could be sequenced, each coming from a transcriptome with unique expression profiles, and analyzed together or individually. If discrete parameter sets are unable to meet the characteristics of desired expression profiles, parameters expressing particular numbers of genes or accumulating particular numbers of transcripts can be decided upon and then a subset of those retained in a modified PRO file to feed back to Flux Simulator for simulated library prep and sequencing.

Rather than subsetting the simulated transcriptome, subsets of the genome annotation itself can be extracted to allow for the expression of sets of genes to be simulated together or for the expression of genes with similar properties to be simulated together. Because the average length of transcripts accumulated to a rare level tends to be longer than the average mRNA length [90], Flux Simulator could be used to simulate partial transcriptomes, one with longer than average transcripts accumulating to the lowest levels and additional partial transcriptomes with other features, that are then merged together for downstream sequencing simulations.

The goal throughout this set of scenarios is to enable the development of methods based on simulated RNA-Seq experiments to test the performance of assembly or other *de novo* methods on particular aspects. Additional simulation tools allow for ways to address a particular problem, while maintaining underlying structure in simulated transcriptomes.

### 3. Simulate transcriptomes for input into other read simulators

Of the RNA-Seq read simulators summarized (Table 1), only Flux Simulator bases transcriptome simulation on Zipf’s law. Therefore it may be desirable to use Flux Simulator’s gene expression output as input into another RNA-Seq read simulator, like Polyester, TuxSim, or RSEM, that can accept user-specified abundances. While RSEM and TuxSim require gene expression data, like that produced by a Flux Simulator-generated transcriptome, Polyester requires read counts for each gene model. This requires a conversion from number of transcripts accumulated to the number of reads expected to map to the transcript, based on sequencing depth and transcript length. Calculating the number of transcripts accumulating from the expression of each gene on a per million basis and equating to Wagner et al.’s [99] calculation of transcripts per million (TPM) gives

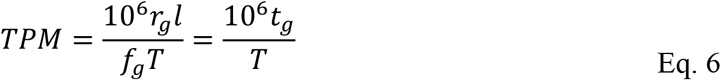

where *r*_*g*_ is number of reads aligning to gene region *g, f*_*g*_ is the length of *g* that has reads mapping to it, *l* is the read length, *T* is the total number of transcripts sequenced. This reduces to

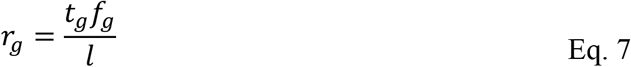

with the number of reads expected to map to gene *g* being the product of the number of transcripts accumulated in the simulation by the expression of the gene *g* and the feature described by the gene model for gene *g* divided by the read length to be sequenced in the sequencing simulation.

This approach can be advantageous when defining a particular model, or set of models, for gene expression but then wanting Polyester’s control over replication or differential expression, RSEM’s learned fragment length distributions or error models, or TuxSim’s options for controlled up and down regulation of transcripts and isoforms. This provides the interface to go from a model of gene expression to read counts.

