## Supplemental Table 1 for "RAFTS: A graphical tool to guide Flux Simulator transcriptome simulation for method development in *de novo* transcriptome assembly from short reads"

Supplemental Table 1. Flux Simulator simulation parameters used across genomes for evaluation, with expectd number of genes expressed.

| Simulation Set | Species inspiring simulation settings | # transcripts in Flux Simulator input | Expected genes expressed, approx | NB_MOLECULES | k | $x_0$ | $x_1$ |
|---|---|---|---|---|---|---|---|
| A | *Brassica napus* | 99,277 | 34,000 | 300,000 | -0.6 | 15,000 | 10,000,000,000 |
| B | *Arabidopsis thaliana* [1] | 25,263 | 10,000 | 100,000 | -0.6 | 5,000 | 100,000,000 |
| C | *Mus musculus* [2] | 35,843 | 8,000 | 3,000,000,000 | -0.6 | 1,000,000 | 5,000,000 |
| D | *Salmo salar* | 85,576 | 25,000 | 60,000,000 | -0.8 | 50,000 | 100,000,000 |
| E | *Saccharomyces cerevisiae* | 5,985 | 4,500 | 300,000 | -0.7 | 50,000 | 5,000,000 |
| F | *Homo sapiens* | 112,892 | 15,500 | 400,000 | -0.9 | 5,000 | 5,000,000,000 |
| G | Flux Simulator defaults | NA | 18,800 | 5,000,000 | -0.6 | 9,500 | 90,250,000 |

[1] Only representative gene models were included in the genome annotation.

[2] Only the genome features annotated as best reference sequences were retained in the genome annotation.

50 replicates: 80% confidence level, margin of error of 20% of the mean for # genes, #SP genes, #CR genes; 80% confidence level, margin of error 33% of the mean for #I genes
125 replicates: 80% confidence level, margin of error of 20% of the mean for # genes, #SP genes, #CR genes, #I genes
300 replicates: 90% confidence level, margin of error of 10% of the mean for # genes, #CR genes; 90% confidence level, margin of error of 25% of the mean for #I genes
1000 replicates: 90% confidence level, margin of error of 10% fo the mean for # genes, #SP genes, #CR genes, #I genes
